# A parasite-inclusive food web for the California rocky intertidal zone

**DOI:** 10.1101/2024.10.14.618335

**Authors:** Zoe L. Zilz, Emily Hascall, Athena DiBartolo, Delén Flores, Sophie Cameron, Jaden E. Orli, Armand M. Kuris

## Abstract

We present a highly resolved, species-rich food web, including parasitic interactions, for the California rocky intertidal zone. The food web is comprised of 1809 nodes, representing 1845 taxa, and 13,222 links representing trophic interactions between nodes. While only 670 links represent parasitic interactions, we have assembled possibly the most speciose parasite-inclusive food web ever published. The inclusion of all nodes and links are justified using multiple lines of evidence which are built into the dataset. In addition, metadata, including trophic strategy, taxonomic information, habitat, and other ecological attributes allow the data user to filter the food web to their specifications. The food web is a powerful and flexible tool for researchers with questions about large network properties, ecological dynamics of rocky shores, and the role of parasites in ecosystems. Our food web can be used to predict how complex ecosystems like the California rocky intertidal will respond to anthropogenic change and management strategies.

## Background & Summary

The California rocky intertidal zone is an iconic habitat with a rich history of foundational ecology and natural history. Due to the biodiversity and ease of access of the rocky intertidal zone (RIZ) ecosystem, it became a hub of experimental ecology in the 1960s and 70s^1,2^; as a result, there is a comprehensive body of literature on RIZ interspecies interactions. Unfortunately, the accessibility and natural appeal that facilitated decades of intertidal research also poses a conservation challenge for the RIZ, especially on California’s heavily populated and developed coastline^3^. Human recreation, visitation, and harvesting erodes the very biodiversity that draws visitors to the RIZ, with variable consequences for the RIZ ecosystem, adjacent beaches, and subtidal reefs^4–9^. Rocky intertidal habitats face multiple stressors in addition to direct anthropogenic disturbance, including sea surface temperature increase^10–12^, sea level rise^13^, ocean acidification^14^, storm intensification^15^, introduced species^16,17^, and disease^18–22^. Contemporary research on the RIZ has demonstrated large scale shifts in species distributions and community-wide response to climate change^23–25^. There is growing evidence that despite apparent long-term stability, the Northeastern Pacific RIZ is on the cusp of an ecological tipping point^26,27^. Especially in extremely biodiverse ecosystems like the RIZ, species losses or shifts in abundance can be cryptic, resulting in subtle effects on ecosystem dynamics that go unnoticed until reversal is too late^28^.

Scientists can use food webs as a tool to predict and mitigate the effects of anthropogenic disturbance. Food webs are valuable tools for contextualizing species interactions, modeling complex community structure, and reconciling data on pairwise relationships with larger-scale ecological patterns^29^. Food webs can be useful to predict ecosystem response to perturbation^30–32^, ecosystem impacts of species loss^33–36^, and can identify key conservation targets^37,38^. Construction of highly resolved food webs, while challenging, can yield powerful tools for conservation, reflecting the true complexity of ecosystems. Ecological network structure (properties of food web networks such as stability, nestedness, centrality, and connectedness) can add depth to simple metrics such as species richness, which has previously been a common focus of conservation efforts^39^. Food webs represent a tangible link between basic research and conservation priorities. For example, loop analysis—an analytical tool demonstrating the effect of a sustained change in abundance of one species on the rest of the food web—can provide critical information to managers as they consider the downstream effects of either protecting or eradicating species^40^. Furthermore, simulating press perturbations using conceptual food webs can help coastal managers identify conservation priorities (e.g. umbrella species) and strategies^41^. In the face of the Anthropocene, developing reliable high-resolution baselines for ecosystems is paramount. In RIZ areas that are difficult to access or are data poor, a highly resolved food web can be a critical tool for making management decisions without site-specific data.

Analytical representations of ecosystems must include their infectious agents. Parasitic organisms form the backbone of essential ecosystem properties and function. These consumers are abundant in other coastal food webs in California^42–44^ and in intertidal habitats worldwide^45–48^. Parasites can alter the way organisms interact with each other, both intra- and interspecifically^49^. Generally, parasites stabilize food webs by increasing connectance and nestedness, although certain parasites’ sensitivity to extinction can negate stabilizing effects^50–52^. Parasite communities can also be indicative of certain ecosystem properties^53^. Many metazoan parasites have complex life cycles that rely on trophic interactions between a sequence of hosts, therefore requiring the presence and abundance of multiple organisms^54^. As a result, the presence or absence of parasites with complex life cycles (trematodes, cestodes, acanthocephalans, nematodes, and some parasitic crustaceans) reflects the population status of (and trophic relationships between) multiple hosts. Furthermore, some trophically transmitted parasites, for example, trematodes and cestodes, require vertebrate hosts for their definitive (sexually reproductive) life stage. Systems experiencing reductions in the abundance of vertebrate top predators may have a paucity of these parasite taxa compared to ecosystems with intact predator communities^55–57^. Characteristics of parasite communities have enabled the use of parasite assemblages in the evaluation of protected areas and of habitat restoration success^45,58–60^.

In the interest of developing a dynamic tool for ecosystem-based management of the California RIZ, we constructed a highly resolved topological food web using the wide body of literature on the RIZ, supplemented with gut content analyses and parasitological assessments for selected host species. Following the framework set forth by authors of other food webs for California’s coastal habitat, primarily the kelp forest food web in Morton *et al* 2021^44^, this web provides a highly taxonomically resolved baseline for the entire California RIZ. Although a geographically comprehensive web cannot represent every local RIZ habitat, this web is designed to be subsetted and manipulated by end users to represent specific rocky intertidal communities of interest. The network includes species from both biogeographic regions (the Oregonian and Californian provinces) present on the California coast. The inclusion of both biogeographic provinces should make the food web a useful starting point for RIZ research extending to the north and south of California. This web comprises 1845 total nodes—1618 of which are resolved to species—including 1443 free-living species (1510 nodes) and 314 parasitic taxa (335 life stages). Among these taxa there are 13,222 trophic links, about 5% of which are parasitic interactions (670 host-parasite links)(Figure 1). The California RIZ food web is perhaps the most speciose parasite-inclusive food web yet constructed. We encourage the use of the food web as a starting point for conducting more complex research on the California rocky intertidal zone ecosystems.

**Figure 1.**
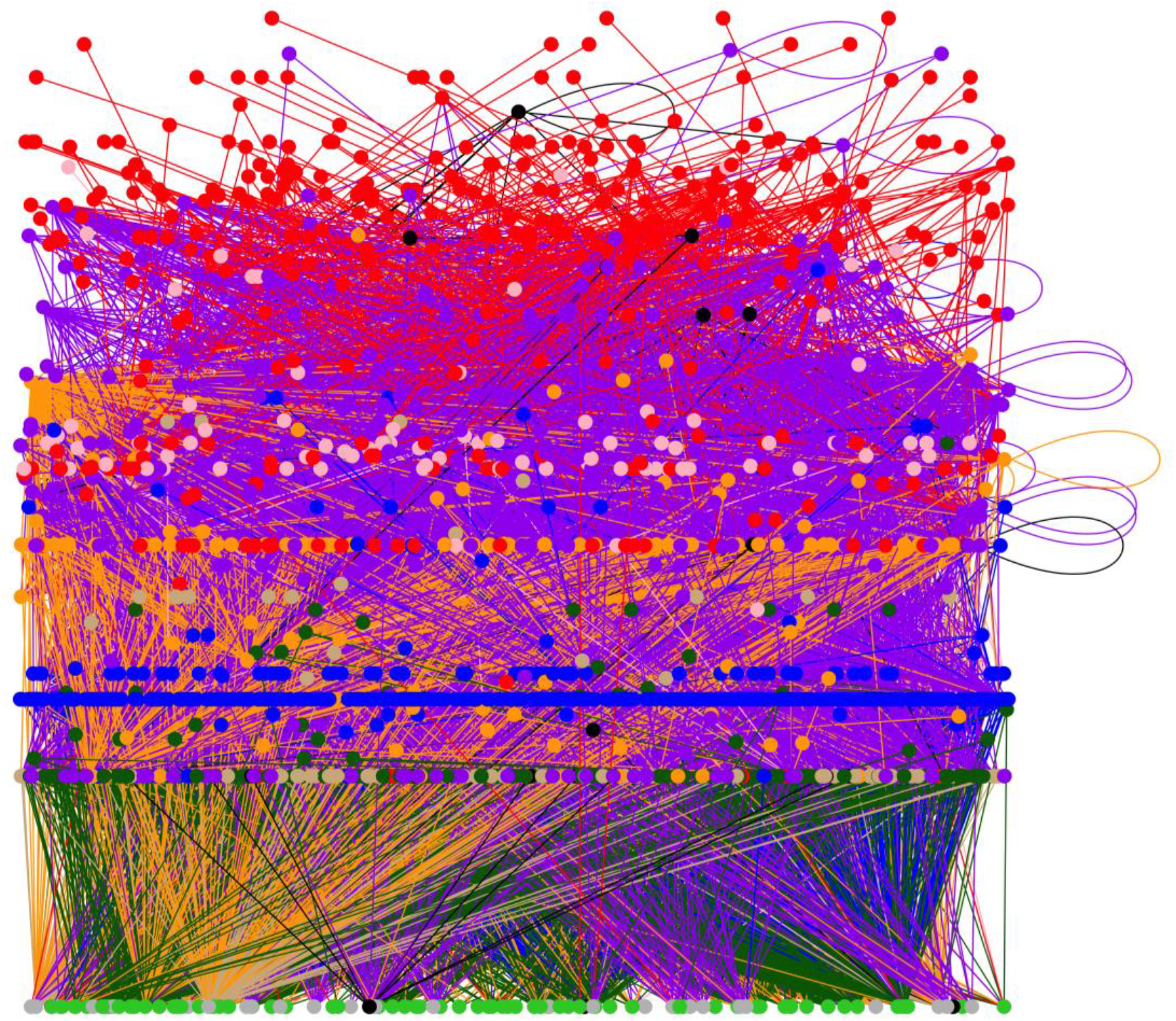
Topographic representation of the food web, inclusive of parasites, for the California rocky intertidal zone. Circles represent nodes, connected by lines to represent links, or interactions between nodes. Loops are indicative of cannibalistic links. Due to the taxonomic richness of the web, several nodes and links overlap, so not all nodes or links are visible. Nodes are organized according to trophic level, with basal nodes (primary producers, non-living material, and non-feeding life stages) at the bottom and top-level consumers at the top. Node colors indicate consumer strategy, as follows: light green = primary producers; grey = non-feeding; brown = scavengers and detritivores; blue = filter feeders; orange = omnivores; purple = carnivore; red = parasites and symbiotic egg predators; black = unknown or various. Link colors are inherited from consumer nodes. An interactive version of this figure is available at https://rpubs.com/zoe_zilz/968403.

## Methods

### Habitat and Study Area Description

Rocky intertidal zones are ecosystems comprised of bedrock or boulders situated between the mean high-water line (supralittoral zone) and the mean lowest low water line (subtidal zone). This zone contains within it several habitats, including the splash zone, high and low tide pools, exposed outcroppings, reef flats, boulder fields, surge channels, surfgrass beds, and mussel clumps.

This study focused on rocky intertidal organisms present along the coast of California, comprised of two major biogeographic zones: the Oregonian Province to the north of Point Conception and the Californian Province to the south. The field sampling component of this research was centered at Santa Barbara, CA, whose proximity to Point Conception results in a mix of southern and northern species (Figure 2). For this reason, we include organisms with geographic ranges both north and south of Point Conception in the meta-food web, allowing end-users to subset the data set to their region of interest.

**Figure 2.**
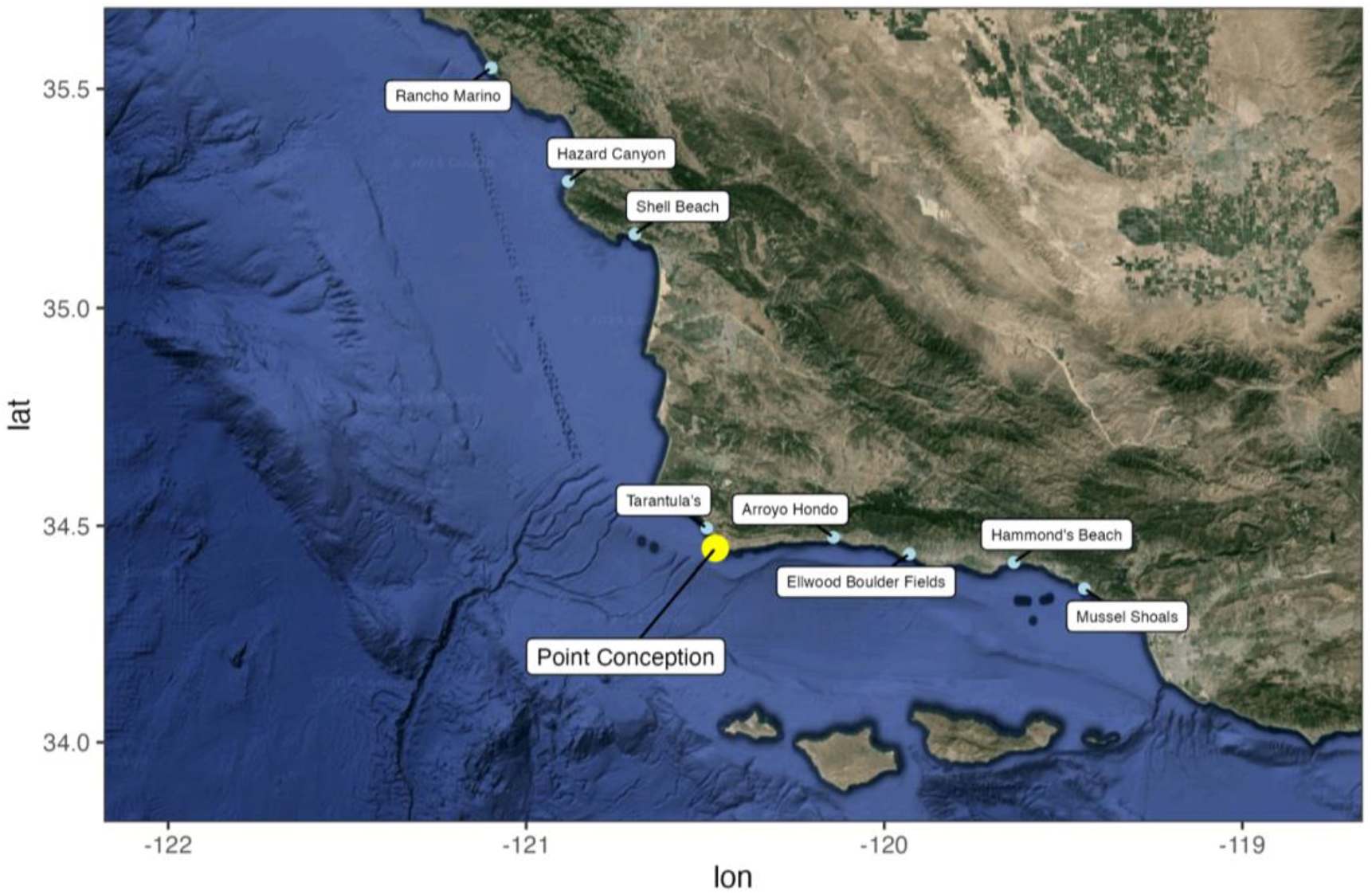
Satellite map of the eight sites used for collection of rocky intertidal zone fish and invertebrates. Point Conception is indicated by a larger yellow circle to signify the biogeographic break between the Oregonian Province (north) and the Californian Province (south).

### Data Sources

In the construction of a food web for the California rocky intertidal we intended to develop a meta-network that represented both published and novel empirical observations. The network is designed to grow and change as new nodes and links are observed. To this end, we used two foundational rocky intertidal guides^61,62^, and expert observers (e.g. University of California Natural Reserve Managers) to first create a comprehensive species list for California. This species list included fishes, invertebrates, microorganisms, algae, and known parasites. The preliminary list was augmented using direct and citizen science observations (e.g. iNaturalist), primary literature, and grey literature.

Birds received special consideration as frequent visitors to the intertidal and important top predators in the system. Unlike for fish and invertebrates, there is no single primary literature source describing rocky intertidal birds, given that many birds are transient or forage additionally in other habitats. Given these challenges, we based our bird list on a systematic survey of birds foraging in intertidal habitats at Coal Oil Point Reserve in Santa Barbara, CA, published in Lafferty (2001)^63^. We were given access to the raw data from the Coal Oil Point Reserve surveys and reanalyzed these data to focus specifically on species actively foraging in the RIZ. We augmented our bird list based on species-specific literature and expert opinion.

Trophic links (including reports of parasitism) were first determined using a systematic literature review (Web of Science and Google Scholar). One major source of interaction data was another comprehensive food web for the Northeastern Pacific rocky intertidal, the Sanak Archipelago, Alaska food web first published by Wood *et al*., 2015^64^, and refined by Dunne *et al*., 2016^30^. We first subsetted the Sanak intertidal food web from Dunne *et al*., 2016^30^, to only include organisms from our California RIZ node list and then reconstructed the interactions using Sanak food web data. We acknowledge that other food webs for the Northeastern Pacific rocky intertidal exist (e.g. Wootton, Sander, and Allesina, 2016^65^), but found that the Sanak food web to be the only one accurate enough, taxonomically resolved enough, and similar enough to our system to justify incorporation. We also conducted targeted sampling of mostly decapod crustaceans and fish on the southern and central coast of California to further determine trophic and parasitic relationships amongst major consumer groups. See Food Web Assembly below for exact food web construction methods.

### Sampling Methods

We collected fishes and invertebrates to include diet and parasitological information for organisms that were underrepresented in the literature or whose trophic relationships were understudied. Additional species were collected opportunistically. Collection sites included eight rocky intertidal zone areas both north (Rancho Marino University of California Natural Reserve, Hazard Canyon, Shell Beach, Jalama County Beach or “Tarantula’s”) and south (Arroyo Hondo, Ellwood Boulder Fields, Hammond’s Beach, Mussel Shoals) of Point Conception, CA (Figure 2).

#### Fishes

Fishes were collected from tide pools and channels by hand using dip nets. Fishes were collected opportunistically and no specific species were targeted, as tide pool fish diets and host-parasite relationships are relatively well studied^66^. Specimens were collected and euthanized in the field under CDFW Scientific Collecting Permit S-190930002-21064-001 and University of California Santa Barbara IACUC protocols 935.1 and 945.1.

#### Invertebrates

Invertebrate species with no known reports of parasites were prioritized for collection. Invertebrate taxa were prioritized that are considered well-suited intermediate hosts for trophically transmitted parasites (e.g. mollusks and crustaceans). Invertebrates were collected by hand opportunistically from tidepools, channels, rocks, crevices, and sediment. Invertebrates were kept alive in flow-through seawater until dissection or frozen for subsequent parasitological examination.

#### Parasitological Assessment

Fishes and invertebrates were thoroughly assessed for potential parasites and diet determination. Exterior tissues (skin or carapace, fins or limbs, gills, mouth, and nasal cavities) were examined under a stereomicroscope for ectoparasites. We used a glass slide to scrape mucus to remove and detect cryptic ectoparasites. All internal organs and musculature were placed between glass plates and “squashed” to create a two-dimensional, relatively transparent field of view which was then scanned under a stereomicroscope for internal parasites. Small (< 20 mm) or soft-bodied invertebrates were squashed whole. We photographed and identified gut contents to the lowest recognizable taxon. Parasites were quantified, photographed, preserved in 80% ethanol, and identified to the lowest recognizable taxon. When species-level identification was not possible, parasites were assigned a morphospecies identification code.

### Food Web Assembly

We closely followed the methods for network assembly outlined by Morton et al., 2021^44^. Our node list was constructed with the intention to expand on the node list generated by Morton et al., 2021, which describes the adjacent kelp forest ecosystem and includes some overlapping taxa. A node present in both the California RIZ and the Santa Barbara kelp forest will only be differently named between the two datasets if the taxonomy of that node has changed between 2021 and the time of this publication; we have included taxonomic synonymies to aid in merging in this rare case. We also use the same life stage, justification, and confidence codes as the dataset in Morton et al., 2021. There are, however, some stylistic differences in the two datasets. For interested readers, we have generated sample R code to fuse the RIZ and kelp forest node lists, which can be found in the Dryad repository associated with this manuscript^67^.

#### Nodes

We systematically justified the inclusion of each node (via literature, expert opinion, or personal observation) in the web and included justification codes and associated references in the data set. We included organisms reported to primarily live in or feed in the RIZ as well as species that regularly visit the RIZ to feed but who are not dependent on the habitat (e.g. birds, kelp-forest or sandy-beach fishes, marine mammals, coastal terrestrial animals). Rare species, as defined by the literature used to justify their inclusion, were excluded. We also explicitly excluded intertidal species that were obligate residents of bays, marinas, fouling communities, soft-bottomed habitats, and subtidal habitats. We did, however, include species from seagrass (*Phyllospadix* sp.) beds since these are often interspersed within rocky areas (author pers. obvs.). Most nodes were resolved to the species level. Regardless of taxonomic resolution, all nodes are accompanied by a permanent identifier (“AphiaID”) that corresponds to an entry in the World Register of Marine Species (WoRMS) database in case of future changes to that node’s taxonomy. Life stages within a species were included as separate nodes when an organism switched trophic function from one life stage to another and these functionally distinct life stages existed in the RIZ. For example, many species of pycnogonids (sea spiders) have parasitic larvae that form galls within their cnidarian hosts but are commensal filter feeders as adults; these pycnogonids therefore were split into a “larva” node and “adult” node to reflect the different trophic functions of each life stage. However, most free-living invertebrates and fish in the RIZ spawn pelagic eggs and larvae that do not remain in the RIZ; these not included as separate nodes unless there was evidence that they stayed in the RIZ (e.g. crawl away larvae of some chitons or egg clusters attached to the benthos by nudibranchs). While we resolved nodes to species wherever possible, some nodes represent either taxonomic or trophic groups. For example, some rocky intertidal zone mollusks graze on a matrix of diatoms on the substrate. To reflect that consumer-resource relationship, we recognized an aggregate node for “benthic diatoms”. In the rare case that the diet of an organism was never reported in the literature and not resolved during our diet assessments, we included an “unknown” node to indicate species critically in need of further study.

#### Links

We performed a systematic literature review of trophic interactions for all nodes using Google Scholar and Web of Science with standard search terms (Genus + species + parasit* and Genus + species + diet*, expanded to Genus + parasit* and Genus + diet if necessary). We also inferred trophic links using logic where appropriate, taking into consideration body size ratios, diet specificity, biogeographic range, and habitat overlap. We also used the presence of trophically transmitted parasites in potential consumer and resource pairs to infer trophic relationships. With respect to parasitism, we only included links that were directly observed or specifically reported in the literature (i.e. we did not infer any host-parasite relationships based on parasite life cycle as in Morton et al., 2021^44^).

Justification codes represent the evidence supporting the inclusion of each link. Often, trophic relationships were both observed directly and reported in the literature; in this case, we coded the link with the strongest possible justification and include all relevant sources. We used node justifications and other metadata to assign confidence levels to each link—ranging from 1 to 5—and included all relevant references. Links where both consumer and resource are known from the California RIZ, and where the interaction between the two was reported or directly observed in the California RIZ, were assigned a confidence level of 1 (very certain). When both species are known from the California RIZ, but interactions between them are only reported outside of California or outside of the RIZ (e.g. bays, subtidal habitats), we assigned a confidence level of 2 (certain). When only one species in the consumer-resource pair is known from the California RIZ and the interaction is only reported outside of California or outside of the RIZ, we assigned a confidence level of 3 (somewhat certain). Consumer-resource links where the consumer is outside of the California RIZ and the trophic interaction has been inferred (see above) we assigned a confidence level of 4 (plausible). We also assigned a confidence level of 4 to inferred interactions where the resource occurs in the same foraging habitat as the consumer and falls within the size range and diet preferences for the consumer. On occasion, organisms in the California RIZ live in close association but the specific nature of their trophic relationship is unknown. For example, many pea crabs (genus *Pinnixia*) live within the mantle cavity of molluscan hosts, but it is unclear whether the relationship is commensal, kleptoparasitic (stealing food from host), or truly parasitic. In these cases we assigned a confidence level of 5 (interaction certain but nature unknown). Literature sources often report diet items at higher taxonomic levels than species. We included these reported interactions with the justification code 6, meaning “more broadly in literature, e.g. group listed but not that species”. The confidence levels for these interactions (where either the consumer or resource was not resolved fully to species) were downgraded by one level. For example, if herring gull (*Larus argentatus*) were reported to eat fish in the genus *Sebastes* in California tide pools, then we would include a link between *L. argentatus* and all *Sebastes* species in our node list, but with a confidence level of 2 instead of 1.

#### Metadata

Additional metadata for each node includes species consumer type (e.g. predator, detritivore, autotroph, parasite, etc. defined by Lafferty et al. (2015), commonality, size range, taxonomic information (phylum, class, order, family), tidal zone association, habitat association (e.g. algae holdfast, water column, rock surface, host), and geographic range. Commonality for most organisms was copied from the primary literature source(s) related to each node. For birds specifically, commonality was determined primarily using raw data on density of birds *actively feeding* in the RIZ at Coal Oil Point Reserve in Santa Barbara, CA, from 1999-2000 and published in Lafferty (2001)^63^ (for RIZ birds not present in these data, commonality was determined as for non-bird species). Metadata for each link includes the interaction type (expanding on the framework defined by Lafferty and Kuris, 2002^69^), rocky intertidal zone where the interaction takes place, the habitat where the interaction takes place (e.g. pools, exposed rock, or both), the locality or localities where the interaction was observed, and references. See column_descriptors.csv for detailed definitions of each metadata type and possible values. This food web is topographic and does not include interaction strength or link weights.

#### Missing and Excluded Links

We acknowledge that many RIZ organisms broadcast spawn and have pelagic offspring, and that the consumption of these propagules by indiscriminate filter feeders is likely an important process within the RIZ foodweb. However, we chose to exclude these assumed feeding interactions from the list of links unless there was explicit evidence of the interaction in our observations or in the literature.

We also acknowledge that, as with other food webs, there are likely many missing links in this web and that those missing links likely represent biases in the greater RIZ research lexicon. For example, our dataset contains relatively few parasitic links compared to other similar food webs^42,44^, which could simply indicate that the RIZ has not been as well sampled for parasitic taxa. However, we are only including links that have referenceable evidence for their inclusion instead of attempting to mathematically estimate the likelihood of real but unobserved links.

#### Food Web Construction

The food web was constructed and visualized using the igraph, cheddar, NetIndices, and VisNetwork packages in R Version 4.4.0^70–73^ (Figure 1).

### Data Records

The California rocky intertidal food web dataset is available from the Dryad digital repository: https://doi.org/10.5061/dryad.3ffbg79sb^67^. The full California rocky intertidal food web dataset includes a node list with metadata (nodes.csv), a link list with metadata (links.csv), a list of references used for assembling the web (references.csv), and a file containing column header and code definitions (column_descriptors.csv).

### Technical Validation

The inclusion of each node and link in the food web was validated using multiple lines of evidence. Nodes and links are accompanied by all references related to their inclusion, even when interactions were inferred using logic. Codes indicating the type of inference (e.g. “inferred based on shared habitat and diet category”) are included for all links. This methodology for assembling food webs has precedence in Morton et al., 2021^44^.

### Usage Notes

The food web dataset is formatted for specifically for use with the R package *cheddar*^71^ but can easily be adapted for use with other analytical tools. Metadata in both the link and nodes data frames allow the food web to be filtered to the end user’s specifications. Nodes can be filtered or aggregated by trophic strategy, rocky intertidal zone, several levels of taxonomy, and/or life stage. Links can be filtered by justification, confidence, or interaction type, and parasitic interactions can easily be added to or removed from the web. The flexibility of the dataset allows for a myriad of analyses by other researchers. For suggestions on how to get started with this dataset, see Code Availability. End users who wish to simply visualize the food web can access an interactive version online at https://rpubs.com/zoe_zilz/968403.

## Code Availability

Annotated code for assembling the food web and generating network figures is available on our Dryad digital repository (https://doi.org/10.5061/dryad.3ffbg79sb) in the cariz_foodweb_visualization.Rmd file. We have also included some suggested code for subsetting the food web network in the example_web_subset_code.Rmd file.

## Acknowledgements

This research was carried out on unceded indigenous land originally occupied by the Chumash, Barbareño Chumash, Obispeño, and ‘Amuwu peoples. Our work was financially supported by the UC Santa Barbara Worster Award and the UC Santa Barbara Coastal Fund. This preparation of this food web and the associated data would not have been possible without a small army of undergraduate researchers, including: Jade Morris, Zoe Fung, Lizzie Wu, Maddie Elkin, Rosa Garcia Alfaro, Maritza Barrena, Kate Nickel, and Elle Allesandrini. In addition, Marisa Morse, Caroline Owens, Sam Bogan, Sriram Ramamurthy, Avrey Parsons-Field, Christoph Pierre, and Amelia Ritger helped with field collections. Kevin Lafferty provided guidance on the early field work, field site selection, and data collection protocols. University of California Natural Reserve System Reserve Managers Cristina Sandoval and Keith Seydel provided access to intertidal areas and invaluable local knowledge. John McLaughlin and Dana Morton provided significant support with project design, reviewing of protocols, and food web assembly. Chelsea Wood and Kevin Lafferty provided unpublished or raw data that was incorporated into the food web.

## Author contributions

Z.L.Z. and A.M.K. designed the study. Z.L.Z., E.H., A.D., D. F., and J.E.O. performed the systematic literature search, collected live specimens, and performed necropsies. S.C. provided significant expertise on the diet and trophic position of rocky intertidal birds. Z.L.Z. assembled the food web.

Z.L.Z. and A.M.K. wrote the manuscript.

All authors read, copy-edited, and approved the manuscript.

### Competing interests

The authors declare no competing interests.

## Notes

https://doi.org/10.5061/dryad.3ffbg79sb

### Competing Interest Statement

The authors have declared no competing interest.

